# Genome of a novel Sediminibacterium discovered in association with two species of freshwater stream Cyanobacteria in Southern California

**DOI:** 10.1101/2021.08.20.457134

**Authors:** David Castro, Ciara Sanders, Sandy Lastor, Andrea Moron-Solano, Briana Vega, Hannah Hausknecht-Buss, Simone Henry, Andrew Zhang, Haven Johansen, Lakme Caceres, Antolette Kasler, Isabelle Massaro, Immanuel Mekuria, Gretchen Vengerova, Rosalina Stancheva Hristova, Xiaoyu Zhang, Betsy Read, Arun Sethuraman

## Abstract

Here we report the discovery of a novel Sediminibacterium sequenced from laboratory cultures of freshwater stream cyanobacteria from sites in Southern California, grown in BG11 media.

## INTRODUCTION

The phycosphere, or the area immediately surrounding cyanobacterial cells or colonies(1), is a resource-rich environment for heterotrophic bacterial colonizers. These colonizers often evolve mutualistically with their host cyanobacteria to provide nutrients and facilitate remineralization of organic material in the environment. The bacterial-cyanobacterial interactions in the phycosphere of lake planktonic cyanobacteria are relatively well studied(2),(3), in contrast to stream benthic cyanobacterial mats. The mucilage of freshwater cyanobacteria and other algae represents a unique habitat and nutrient source, which is beneficial for heterotrophic bacteria(3) and some mixotrophic endogloeic diatoms(4). Here we report the discovery of a novel Sediminibacterium sequenced from laboratory cultures of freshwater stream cyanobacteria from sites in Southern California, grown in BG11 media. A de novo genome assembly of this bacterium yielded a complete chromosomal genome on a single contig of length 3.34 Mbps with a GC content of 39.4%. A first pass annotation identified 3000 protein coding genes, with 98% completeness when compared against all prokaryotic and cyanobacterial gene families in BUSCO. A comprehensive phylogenomic species tree reconstruction using 100 of these protein coding genes placed the novel bacterium to be sister to a previously identified Sediminibacterium, which is over 20% divergent from our novel genome.

## PROVENANCE

The novel bacterium was isolated from cultures of two separate cyanobacterial species, morphologically identified as Chamaesiphon (epilithic, epiphytic) and Lyngbya-like (benthic), sampled from Escondido Creek in December 2018 and Anza Borrego Desert stream in December 2019. Escondido Creek is an urban stream subject to intense agricultural runoff, increased salinity and nutrients. The cleaner, intermittent creek in Anza Borrego was nearly completely dry during sampling. Both samples were returned to the California Algae Lab in CSU San Marcos, CA for cyanobacteria isolation. Chamaesiphon cells were initially cultured on a solid BG-11 medium until dispersed colonies developed and these were used for monoclonal strain isolation and growth in liquid BG-11, while a single filament of Lyngbya-like was isolated directly in liquid BG-11. Both non axenic cyanobacterial strains grew for 50 days at 20-23°C with an irradiance of 80mmol photons m^−2^s^−2^ and a 12:12h light:dark cycle(5). Transmission electron microscopy was performed with fresh cultured material from both cyanobacterial strains.

## METHODS

DNA extraction was performed using the CTAB chloroform extraction protocol and uniquely barcoded whole genome libraries were prepared using Transposase Enzyme Linked Long-read Sequencing (TELL-Seq™) Technology (Universal Sequencing, Carlsbad, CA) using their whole genome sequencing library prep kit. The libraries were pooled, and paired-end sequenced using TC NextSeq™ 500/550 Mid Output Kit V2.5 (150 cycles). Demultiplexed raw reads were subjected to quality control using FastQC(6), and bases with a PHRED Q score of < 30 were trimmed. Raw reads were then assembled using the TELL-Seq™ software. Quality and completeness of the assembly was assessed using QUAST v. 5.0.2(7) and BUSCO v. 5.0.0(8) against the bacteria-odb10 gene family database followed by gene calling and functional analyses using the NCBI Prokaryotic Genome Annotation Pipeline (PGAP(9)) and KEGG(10). Annotated protein coding sequences were then used as queries against the nr database in NCBI using BLASTP(11), and the top 50 hits were obtained as multiple sequence alignments with Mview(12). RAxML(13) was then used to generate gene trees and a random set of 100 gene trees were pooled to generate a consensus species tree using ASTRAL(14). A BLASTN homology search was also performed to compare and establish the novelty of our genome against sister species of Sediminibacteria identified in the ASTRAL species tree.

## RESULTS

Chamaesiphon colonies consisted of numerous ovoid cells surrounded by individual mucilagenous sheath and embedded in a common gelatinous matrix, while the cells of Lyngbya-like cyanobacterium were arranged in filaments enclosed in a thick multilayered sheath. Oval to cylindrical bacterial cells, presumably belonging to Sediminibacteria, were distributed inside the common gelatinous matrix of Chamaesiphon colony (Fig:1A) and on the top of the mucilagenous sheath of Lyngbya-like filaments.

**FIG 1.**
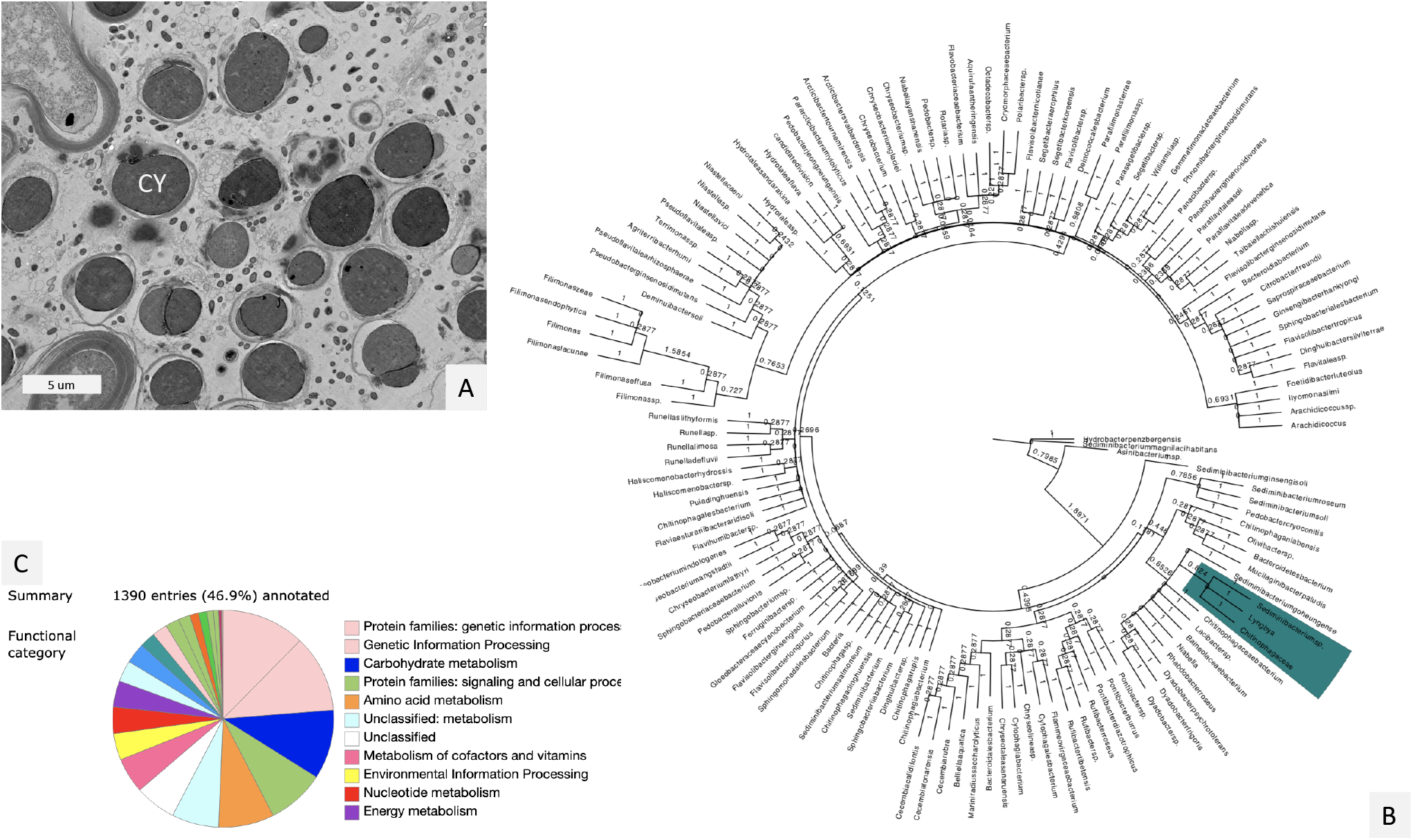
Fig. 1A is a TEM micrograph showing bacteria, possibly our novel Sediminibacterium within the phycosphere of the cultured Chamaesiphon (CY). Fig. 1B shows a phylogenomic species tree analysis which with the polytomic ancestry of our novel Sediminibacterium marked in color. Fig. 1C shows the KEGG functional classification of approximately 47% of all annotated genes.

2.1 x 10^9^ raw reads were sequenced prior to assembly.Genome assemblies from both sites revealed the presence of a single species of bacterium (99% identity between assemblies). Here we report the most contiguous of the assemblies (1 contig, 3.34 Mbps, 39.4% GC), which has been submitted to NCBI (Accession ID:). The novelty of this genome was established using genomic similarity with 28 other Sediminibacterial genomes, with the highest matches (81.46%) to NCBI Accession IDs: GCA013391385.1, GCA019264545.1, and GCA019264645.1. Annotation using PGAP identified 3009 protein coding genes, which included 45 RNA sequences (3 rRNA, 39 tRNA, 3 ncRNA). The genome was identified to be 97% complete (BUSCO, bacterial-odb10), with 1% of duplications, and 2% of missing families. 47% of all annotated genes were functionally classified using KEGG (Fig:1B). Species tree reconstruction using ASTRAL placed our novel bacterium as sister to Sediminibacteria, with an unresolved polytomy with hot-spring Chitinobacteria within the same super-clade (Fig:1C).

## ACKNOWLEDGMENTS

This work was funded by NSF-REU: 1852189 to PI Betsy Read and co-PI Sethuraman, and was conducted over Summer 2021 by CSUSM NSF REU students whose experiences were chronicled at www.csusmbioreu.weebly.com.

## REFERENCES

1. Ramanan R, Kim BH, Cho DH, Oh HM, Kim HS. 2016. Algae-bacteria interactions: evolution, ecology and emerging applications. Biotechnol advances 34(1):14–29.

2. Woodhouse JN, Ziegler J, Grossart HP, Neilan BA. 2018. Cyanobacterial community composition and bacteria-bacteria interactions promote the stable occurrence of particle-associated bacteria. Front microbiology 9:777.

3. Brunberg AK. 1999. Contribution of bacteria in the mucilage of Microcystis spp.(Cyanobacteria)to benthic and pelagic bacterial production in a hypereutrophic lake. FEMS Microbiol Ecol 29 (1):13–22.

4. Stancheva R, Lowe R, Lowe R. 2019. Diatom symbioses with other photoautotroph. Diatoms: Fundam Appl Seckbach, J., Gordon, R., Eds p 225–244.

5. Conklin KY, Stancheva R, Otten TG, Fadness R, Boyer GL, Read B, Zhang X, Sheath RG. 2020. Molecular and morphological characterization of a novel dihydroanatoxin-a producing Microcoleus species (cyanobacteria) from the Russian River, California, USA. Harmful algae 93:101767.

6. FastQC. 2016. FastQC: a quality control tool for high throughput sequence data..

7. Gurevich A, Saveliev V, Vyahhi N, Tesler G. 2013. QUAST: quality assessment tool for genome assemblies. Bioinformatics 29 (8):1072–1075.

8. Seppey M, Manni M, Zdobnov EM. 2019. BUSCO: assessing genome assembly and annotation completeness. Methods molecular biology (Clifton, NJ) 1962:227–245.

9. Tatusova T, DiCuccio M, Badretdin A, Chetvernin V, Nawrocki EP, Zaslavsky L, Lomsadze A, Pruitt KD, Borodovsky M, Ostell J. 2016. NCBI prokaryotic genome annotation pipeline. Nucleicacids research 44 (14):6614–6624.

10. Kanehisa M, Araki M, Goto S, Hattori M, Hirakawa M, Itoh M, Katayama T, Kawashima S, Okuda S, Tokimatsu T, et al. 2007. KEGG for linking genomes to life and the environment. Nucleic acids research 36 (suppl_1):D480–D484.

11. Altschul SF, Gish W, Miller W, Myers EW, Lipman DJ. 1990. Basic local alignment search tool. J molecular biology 215 (3):403–410.

12. Brown NP, Leroy C, Sander C. 1998. MView: a web-compatible database search or multiple alignment viewer. Bioinform (Oxford, England) 14 (4):380–381.

13. Stamatakis A. 2014. RAxML version 8: a tool for phylogenetic analysis and post-analysis of large phylogenies. Bioinformatics 30 (9):1312–1313.

14. Mirarab S, Reaz R, Bayzid MS, Zimmermann T, Swenson MS, Warnow T. 2014. ASTRAL: genome-scale coalescent-based species tree estimation. Bioinformatics 30 (17):i541–i548.

15. Willrodt D. 2012. genomediff user manual..

16. Stanke M, Keller O, Gunduz I, Hayes A, Waack S, Morgenstern B. 2006. AUGUSTUS: ab initio prediction of alternative transcripts. Nucleic acids research 34 (suppl_2):W435–W439.

